# Impact of Erg11 amino acid substitutions identified in *Candida auris* clade III isolates on triazole drug susceptibility

**DOI:** 10.1101/2021.08.16.456589

**Authors:** Benjamin Williamson, Adam Wilk, Kevin D. Guerrero, Timothy D. Mikulski, Tony N. Elias, Indira Sawh, Geselle Cancino-Prado, Dianne Gardam, Christopher H. Heath, Nelesh P. Govender, David S. Perlin, Milena Kordalewska, Kelley R. Healey

## Abstract

*ERG11* sequencing of 28 *Candida auris* clade III isolates revealed the presence of concomitant V125A and F126L substitutions. Heterologous expression of Erg11-V125A/F126L in *Saccharomyces cerevisiae* led to reduced fluconazole and voriconazole susceptibilities. Generation of single substitution gene variants through site-directed mutagenesis uncovered that F126L primarily contributes to the elevated triazole MICs. A similar, yet diminished pattern of reduced susceptibility was observed with long-tailed triazoles posaconazole and itraconazole for V125A/F126L, F126L, Y132F, and K143R alleles.

## Text

*Candida auris* is an emerging fungal pathogen that has spread across the globe and caused multiple healthcare center outbreaks. Strains of *C. auris* are divided into five genetically-distinct, geographic clades: South Asian (I), East Asian (II), African (III), South American (IV), and Iranian (V) (1). Initial spread of *C. auris* to the U.S. and other parts of the world is predicted to have occurred through multiple travel-related introductions (2). Recently, several reports have shown high rates of *C. auris* candidemia in hospitalized patients with severe COVID-19 (SARS-CoV-2 virus) infection, particularly in severely ill patients in the ICU setting (3-5). Interestingly, the pathogenicity of *C. auris* differs from other species in that it can colonize the skin, persist on hospital surfaces and medical equipment, and transfer from person to person (6, 7). In addition, *C. auris* exhibits elevated rates of antifungal resistance. Clinical isolates that demonstrate reduced susceptibility to one or more classes of antifungals, including triazoles, polyenes (amphotericin B), and echinocandins, have been reported, with triazole resistance being the most prevalent (8-10).

Triazole antifungals (e.g. fluconazole, voriconazole, itraconazole, posaconazole) target the biosynthesis of fungal ergosterol, specifically through inhibition of lanosterol 14-alpha-demethylase (Erg11p) that is encoded by the *ERG11* gene in yeast. Early reports identified single Erg11 substitutions (F126L, Y132F, or K143R) in strains from multiple clades (9, 11-13). These substitutions were highlighted due to their connection to triazole resistance within other species of *Candida*, specifically *C. albicans* (14). In our previous study (15), we identified and analyzed *C. auris ERG11* mutations in clinical isolates of clade I and IV. Using a heterologous expression system, we directly linked Y132F and K143R Erg11 substitutions to fluconazole and voriconazole resistance; whereas, other alterations (e.g. I466M, Y501H, and clade-specific polymorphisms) were not associated with elevated minimum inhibitory concentrations (MICs).

Here, we investigated triazole resistance in 28 clinical isolates of *C. auris* clade III obtained from South Africa (n=21), Australia (n=5), and the CDC and FDA Antimicrobial Resistance (AR) Isolate Bank (n=2). *ERG11* was amplified and sequenced as described before (15). In agreement with recent reports (8, 16, 17), we identified two *ERG11* mutations, T374C and T376C, that lead to two amino acid substitutions, V125A and F126L, respectively, in all 28 isolates (**Table 1**). Antifungal susceptibility testing was performed according to CLSI methodology (18, 19) and MICs were interpreted using tentative breakpoints as suggested by the CDC (https://www.cdc.gov/fungal/candida-auris/c-auris-antifungal.html). These isolates demonstrated reduced triazole susceptibilities, specifically to fluconazole and voriconazole (**Table 1**). Of note, three clinical isolates (SA 13, 15, and 16) demonstrated fluconazole MICs in the susceptible range (< 32 µg/ml), despite containing the same *ERG11* mutations as the other strains (**Table 1**). This may point to additional alterations in these isolates that specifically influence the fluconazole-Erg11p interaction. Further analyses on these strains are underway.

**Table 1.**
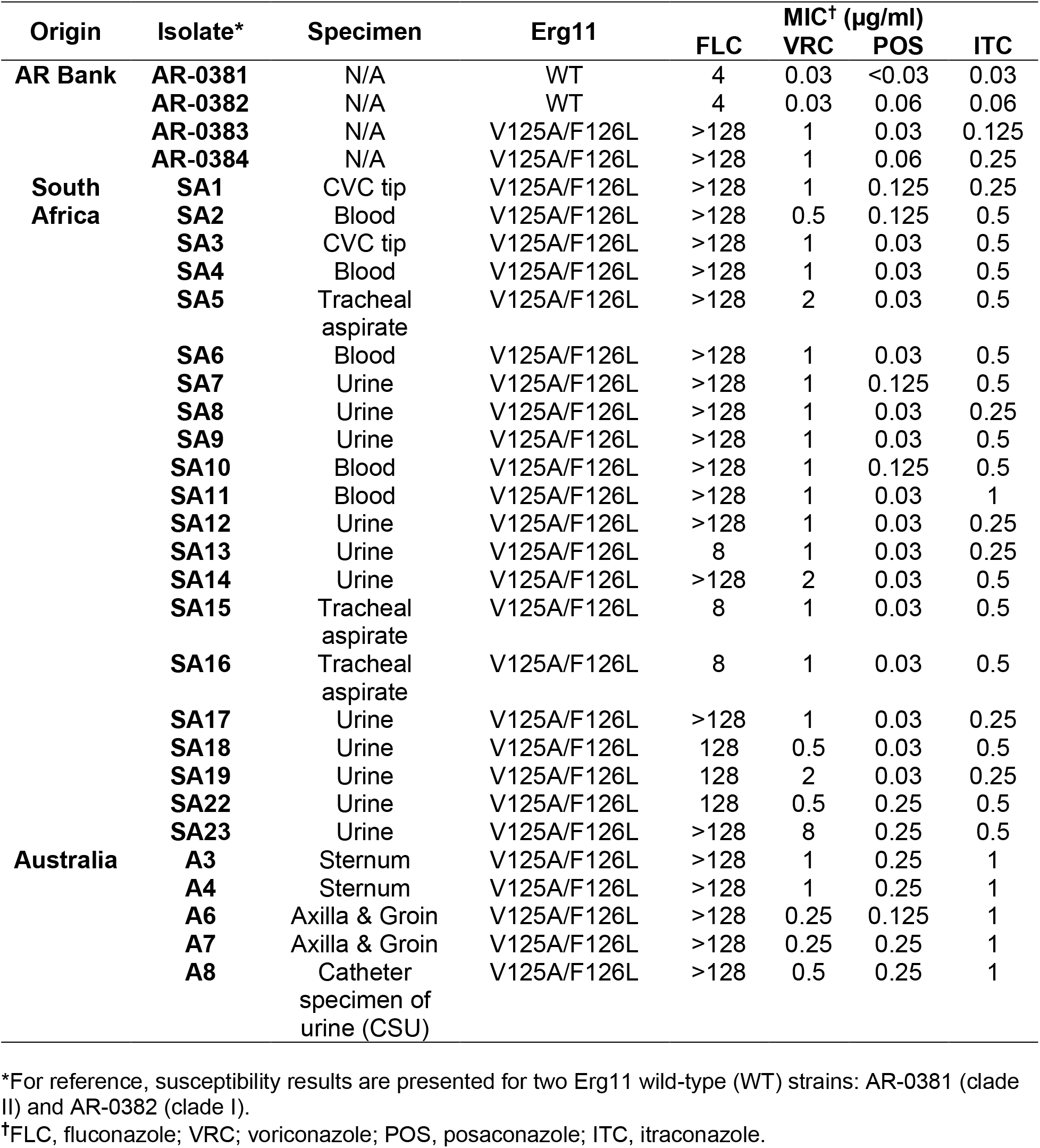
Triazole drug susceptibility and Erg11 profiles of 28 *C. auris* clinical isolates of clade III.

Using the same approach as in (15), we cloned the Erg11 allele (V125A/F126L) from isolate AR-0384 onto a low-copy plasmid (pRS416), which was then expressed in a haploid strain of *S. cerevisiae* (BY4741). This heterologous system allowed us to focus solely on the effects of *ERG11* mutations on triazole susceptibilities. Multiple clones were passaged on selective medium (synthetic defined medium lacking uracil; SD-Ura), screened by PCR, and resulting plasmid sequences verified (for primers, see (15)). *S. cerevisiae* that expressed *C. auris* Erg11-V125A/F126L demonstrated elevated MICs to fluconazole (64 µg/ml) and voriconazole (1 µg/ml). In comparison, expression of an empty vector or Erg11-wild type alleles from other clades yielded MICs 4 to 8-fold more susceptible (≤ 16 µg/ml to fluconazole; ≤ 0.25 µg/ml to voriconazole) (**Table 2**).

**Table 2.** Triazole susceptibilities of *S. cerevisiae* strains that express *C. auris* Erg11 plasmid constructs.

| Erg11 allele | MIC* (µg/ml) |  | POS | ITC |
| --- | --- | --- | --- | --- |
|  | FLC | VRC |  |  |
| Empty vector | 16 | 0.12 | 0.25 | 0.5 |
| wild type (clade I) | 16 | 0.25 | 0.25 | 1 |
| wild type (clade IV) | 16 | 0.12 | 0.25 | 1 |
| I466M | 16 | 0.25 | 0.5 | 2 |
| Y132F | 128 | 2 | 0.5 | 4 |
| K143R | 64 | 1 | 0.5 | 2 |
| V125A/F126L | 64 | 1 | 0.5 | 2 |
| wild type (clade III) | 16 | 0.25 | 0.25 | 1 |
| V125A | 16 | 0.25 | 0.25 | 1 |
| F126L | 64 | 1 | 0.5 | 4 |

\*MICs are representative of three independent experiments and obtained in both nutrient-rich YPD (Yeast extract, Peptone, Dextrose) (MICs displayed) and nutrient-limited SD-Ura broth media; we observed a 2-fold or less difference between media. Fluconazole and voriconazole MICs of the first six strains were previously reported (15); however, these MICs were repeated in tandem with the newly engineered strains and are presented here for comparison. FLC, fluconazole; VRC; voriconazole; POS, posaconazole; ITC, itraconazole.

To further dissect the specific role of V125A and F126L substitutions in triazole resistance, we designed mutagenic primers to individually revert each amino acid substitution (**Figure 1**). A Phusion site-directed mutagenesis kit (Thermo Scientific, MA, USA) was used to introduce the desired wild-type mutations. The resulting *C. auris* Erg11-V125A and Erg11-F126L plasmid constructs were expressed in *S. cerevisiae*. In addition, we performed two consecutive rounds of site-directed mutagenesis to produce a strain that carried neither substitution (Erg11-V125/F126) (**Figure 1D-E**). This strain represented a *de facto* clade III wild-type allele. Plasmid sequences of all alleles were confirmed. Subsequent triazole susceptibility assays revealed that cells expressing F126L alone exhibited elevated MICs similar in levels to V125A/F126L, while V125A alone led to MICs similar to the wild-type alleles (**Table 2**). Our engineered, clade III wild-type allele yielded susceptible MICs, allowing us to conclude that the *ERG11* mutations, as opposed to expression levels, were mainly contributing to the observed decreased susceptibility.

**Figure 1.**
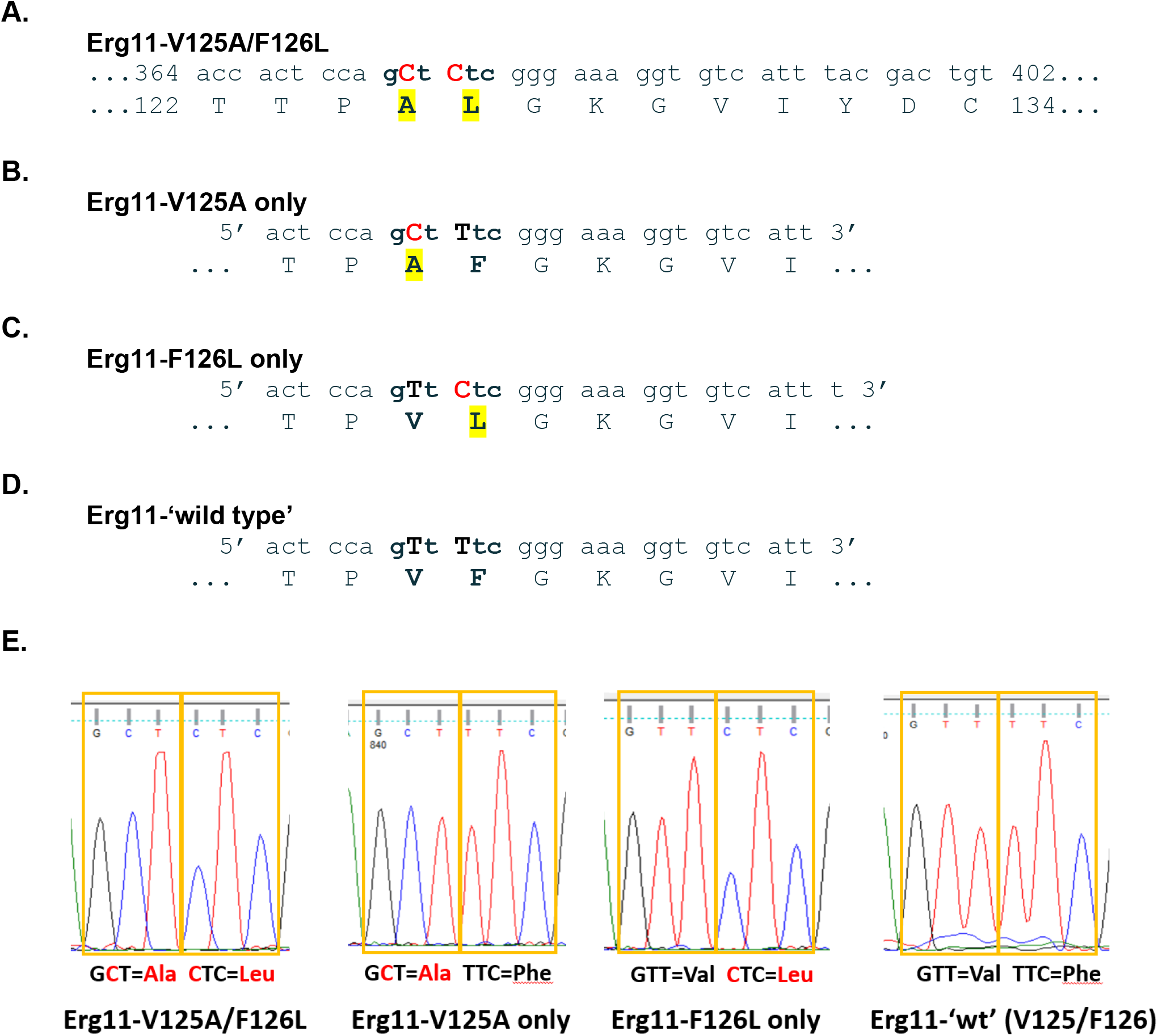
Molecular dissection of *C. auris* Erg11 V125A and F126L amino acid substitutions. **A**, Region of clade III *ERG11* DNA that displays nucleotide mutations (T374C/T376C) in red and resulting protein alterations (V125A/F126L) in yellow highlight. **B**, Forward mutagenic primer used to revert Leucine (L) back to Phenylalanine (F). After mutagenesis, this construct contained only V125A (pCauErg11-V125A). **C**, Forward mutagenic primer used to revert Alanine (A) back to wild type Valine (V). After mutagenesis, this construct contained only F126L (pCauErg11-F126L). **D**, Forward mutagenic primer used to revert Leucine (L) back to Phenylalanine (F) using the pCauErg11-F126L plasmid as a template. After mutagenesis, this construct contained both wild type nucleotides and amino acids (pCauErg11-‘wt’). **E**, Plasmid sequencing chromatograms of relevant codons corresponding to the 125^th^ and 126^th^ amino acids following mutagenesis and propagation in *E. coli*.

Drug binding and cloning studies have demonstrated that certain *ERG11* mutations in *S. cerevisiae* and *C. albicans* influence susceptibility to all triazoles, while other mutations lead to decreased susceptibility to only short- or long-tailed triazoles (20-22). Therefore, in addition to fluconazole and voriconazole (short-tailed triazoles), we tested all of our strains to determine susceptibility to posaconazole and itraconazole (long-tailed triazoles) (**Table 2**). Changes in posaconazole and itraconazole MICs were minimal, although consistent, with 2 to 4-fold differences between the “resistant” alleles (V125A/F126L, Y132F, or K143R) and wild type alleles (**Table 2**). These results are in alignment with the minimal differences observed in clinical isolates (**Table 1**) and to those of previous studies that analyzed Y132F and K143R (or equivalent changes) in *C. albicans* and *S. cerevisiae* (21, 23).

Crystallization of *C. albicans* Erg11 identified residue 126 (and the equivalent residue in *S. cerevisiae*) as being located within the enzyme’s active site and a likely player in substrate binding (24, 25). Furthermore, the authors from this study predicted that alteration of this residue would likely reduce affinity for all triazole drugs but would do so most extensively for short-tailed azoles (24). Since all *C. auris* clade III isolates described in the literature contain both V125A and F126L substitutions, it is likely that these two mutations occurred at nearly the same time in the evolution of this clade. The V125A substitution may simply be a passenger mutation. Alternatively, V125A may increase the stability of the Erg11 enzyme or be advantageous for the yeast in another way and/or in combination with other alterations (e.g., *ERG11* copy number variants (8)). Studies have since identified *TAC1b* transcription factor mutations, linked to increased expression of drug efflux pumps (e.g., *CDR1* and/or other unidentified transporters), as an alternate mechanism of triazole resistance in *C. auris* (26-28). Additionally, a recent report demonstrated an additive effect that concomitant *ERG11* (F444L) and *TAC1b* mutations can have on triazole susceptibility (29).

In conclusion, the *ERG11* allele found in *C. auris* clade III isolates directly contributes to reduced triazole susceptibility, in particular to fluconazole and voriconazole. Moreover, our mutagenic experiments revealed that the F126L substitution was primarily responsible for the elevated triazole MICs. Results of this study further improve our understanding of triazole resistance mechanisms in *C. auris* which can have a direct impact on diagnostic and treatment practices.

## Acknowledgments

This research was supported by William Paterson University (WPU) Department of Biology and College of Science and Health’s Center for Research to K.R.H. Undergraduate work of B.W. and A.W. was supported by WPU College of Science and Health and of I.S. and G.C.-P. by the NSF Louis Stokes Alliances for Minority Participation (LSAMP) program. The authors declare no potential conflicts of interest related to this work.

